# A single-cell method to map higher-order 3D genome organization in thousands of individual cells reveals structural heterogeneity in mouse ES cells

**DOI:** 10.1101/2020.08.11.242081

**Authors:** Mary V. Arrastia, Joanna W. Jachowicz, Noah Ollikainen, Matthew S. Curtis, Charlotte Lai, Sofia A. Quinodoz, David A. Selck, Mitchell Guttman, Rustem F. Ismagilov

## Abstract

In eukaryotes, the nucleus is organized into a three dimensional structure consisting of both local interactions such as those between enhancers and promoters, and long-range higher-order structures such as nuclear bodies. This organization is central to many aspects of nuclear function, including DNA replication, transcription, and cell cycle progression. Nuclear structure intrinsically occurs within single cells; however, measuring such a broad spectrum of 3D DNA interactions on a genome-wide scale and at the single cell level has been a great challenge. To address this, we developed single-cell split-pool recognition of interactions by tag extension (scSPRITE), a new method that enables measurements of genome-wide maps of 3D DNA structure in thousands of individual nuclei. scSPRITE maximizes the number of DNA contacts detected per cell enabling high-resolution genome structure maps within each cells and is easy-to-use and cost-effective. scSPRITE accurately detects chromosome territories, active and inactive compartments, topologically associating domains (TADs), and higher-order structures within single cells. In addition, scSPRITE measures cell-to-cell heterogeneity in genome structure at different levels of resolution and shows that TADs are dynamic units of genome organization that can vary between different cells within a population. scSPRITE will improve our understanding of nuclear architecture and its relationship to nuclear function within an individual nucleus from complex cell types and tissues containing a diverse population of cells.

## INTRODUCTION

In eukaryotes, linear DNA is packaged in a three-dimensional (3D) arrangement within the nucleus. This includes organization of DNA regions from the same chromosome (chromosome territories)^1^ which are further subdivided into megabase-sized self-associating topologically associating domains (TADs)^2,3^, based on gene activity (active/inactive or A/B compartments)^1^, and local interactions between regulatory elements (enhancer-promoter loops)^4–6^. In addition, DNA regions from multiple chromosomes are organized around nuclear bodies that form higher-order structural units^7,8^.

Genome organization within a single nucleus directly impacts various nuclear functions within that cell including DNA replication^9^, transcription^5,10^, and RNA processing^11,12^. Indeed, genome structure is known to dynamically change between different cell types and within individual cells across time to reflect differences in their biological state^5,13,14^. For example, during the cell cycle DNA structure undergoes drastic rearrangement from open chromatin during interphase to highly condensed metaphase chromosomes^15–17^. Similarly, gene expression levels are heterogeneous among populations of cells^18,19^, suggesting that there may be differences in enhancer-promoter contacts present within each cell in the population.

Currently, most methods that are used to study nuclear organization measure ensemble structures across millions of cells. However, because genome structure intrinsically occurs within single cells, ensemble measurements can obscure critical information about the genome organization of any given cell. For example, measuring cells across the cell cycle and averaging their DNA contacts as one population would mask cell-cycle dependent dynamics of the process. Additionally, several studies have shown that observation of genome structures such as TADs^13–16,20^ within single cells do not always match structures predicted from ensemble measurements^1,3,21^. This raises a concern that genome organization observed in bulk assays may not accurately reflect genome structure that exists within biological populations, including structural dynamics or structural heterogeneity.

The two main techniques for measuring genome architecture at the single-cell level are microscopy and single-cell HiC (scHiC). Microscopy provides the capability to study a broad range of genomic interactions in single cells, but is limited to measurements of a small number of loci simultaneously^13,14,20^. In contrast, scHiC provides a genome-wide view of nuclear structure within single cells. However, it is laborious, generates data for few single cells (∼12 cells/experiment), and is limited to low resolution structures (∼10Mb resolution/cell)^15,16,22^. Additionally, neither microscopy nor scHiC can capture long-range interactions (including higher-order structures such as nuclear bodies) across the entire genome.

To address the gap in existing techniques, we developed single-cell split-pool recognition of interactions by tag extension (scSPRITE) as a new, simple and cost-effective method to provide comprehensive, high resolution, genome-wide maps of DNA structure from thousands of single cells. scSPRITE measures both inter- and intrachromosomal interactions and dramatically increases the number of detected DNA contacts per cell relative to existing methods. To demonstrate the utility of scSPRITE, we measured 3D genome structures in >1000 individual mESC nuclei. We observed chromosome territories, A/B compartments, and TADs within hundreds of individual cells. Moreover, we identified higher-order structures in hundreds of single cells, including inter-chromosomal contacts around centromeres, the nucleolus, and nuclear speckles. Interestingly, we identified cell-to-cell heterogeneity in mESC genome structure at different levels of resolution, including at promoter-enhancer contacts of the key pluripotency gene Nanog. These results demonstrate that individual mESC genomes are organized into dynamic and heterogeneous TAD-like units and higher-order structures. Taken together, these observations demonstrate that scSPRITE accurately measures genome structure and uncovers new insights into genome organization. We expect that this approach will enable many future studies examining the relationship between genome organization and nuclear function within individual cells.

## RESULTS

### Single cell SPRITE (scSPRITE): a method to map 3D organization across thousands of single cells

We extended the previously described SPRITE protocol^7^ to map 3D genome structure in thousands of individual single cells. First, we isolate single cells, crosslink DNA and protein complexes, and digest DNA in-nuclei into fragments using restriction enzymes. Then, we perform two sets of split-and-pool barcoding (**Figure 1a**): (i) Nuclear barcoding to tag all DNA molecules contained in a single cell with a unique cell-specific barcode (cell-barcode). In this step, ∼10^5^ nuclei are distributed across a 96-well plate, where each well contains a unique DNA barcode tag such that DNA molecules within the nuclei contained in a well are labeled with the same tag. After ligation of the tags, nuclei are pooled together and the process of split-pool is repeated for a total of three rounds, providing ∼10^6^ unique barcode combinations. Next, we filtered nuclei and sonicated ∼10^3^ nuclei (1% of the total) to produce fragments that we refer to as “spatial complexes”. Although there is no limit in the number of individual cells that can be analyzed using this approach, we focused on ∼10^3^ nuclei to ensure that we achieved high-resolution per cell at a given sequencing depth. (ii) Spatial barcoding uniquely barcodes all DNA molecules within a spatial complex in a given cell. Next, DNA complexes are reverse crosslinked and sequenced. We identify DNA molecules in the same spatial complex by matching all six barcode sequences, and all complexes within the same cell by matching the first three barcodes (**Figure 1a**, see **Methods**)

**Figure 1:**
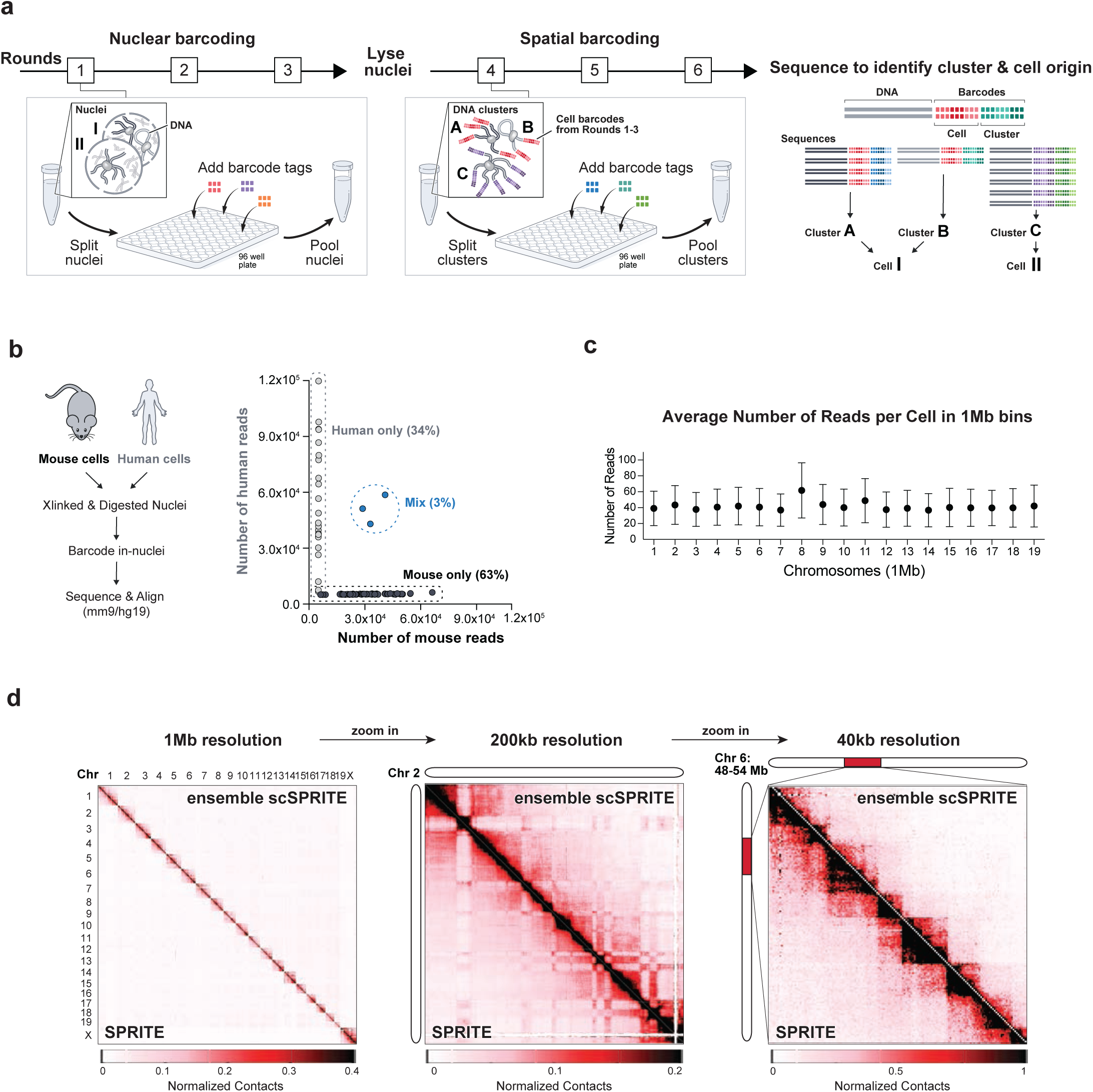
Single cell SPRITE – a novel single cell method to map DNA structure genome-wide. **a**, Schematic of scSPRITE protocol. First, cells are crosslinked, permeabilized, their DNA is digested with restriction enzymes, and nuclei are isolated (see Methods). Second, nuclei are split across a 96-well plate to ligate cell-specific barcodes (each well contains a unique DNA barcode sequence), and pooled together; after three rounds of split-pool nuclei are lysed to release DNA clusters. Third, DNA clusters are split across a 96-well plate to ligate cluster-specific barcodes and pooled together, all for three rounds. Fourth, DNA is sequenced to identify and match barcode sequences of DNA molecules originating from the same cluster and cell. **b**, Estimate of the frequency of cell clumping: equal numbers of mESC and HEKs were combined into the same tube,; crosslinking, in-nuclei digestion, and in-nuclei barcoding were performed. Number of reads for each identified cell barcode ID is plotted. Threshold of >95% single species reads was applied to identify mouse or human only cells. **c**, Average number of reads per single cell in 1Mb bins of all chromosomes; Average and SD shown. **d**, Comparison of scSPRITE (upper diagonal) and SPRITE7 (lower diagonal). Chromosome territories across all chromosomes at 1Mb resolution (left); A/B compartments on chromosome 2 at 200-kb resolution (middle); TADs within a 18-Mb region of chromosome 6 at 40-kb resolution (right).

To validate the single cell approach, we performed scSPRITE on a mixed cell population containing equal numbers of mouse (mESC) and human (HEK) cells pooled together before cross-linking (**Figure 1b**). We split reads into groups based on their cell-barcodes and computed the percentage of reads that aligned exclusively to the mouse or human genome (but not both). We found that >90% of cell-barcode groups contained reads from only a single species, suggesting that the vast majority of cell-barcode groups represent single cells.

We focused our subsequent analyses on mESCs because their genome structure has been extensively studied^3,15^ and they display known functional heterogeneity^17,23,24^. From ∼1500 sequenced mESC nuclei we analytically excluded cell-barcode groups that were likely to represent aggregates by using the number of DNA clusters within cell-barcode groups from a single species (from the previously described mixing experiment). Additionally, to ensure that only intact cells with the highest coverage per cell were analyzed, we focused our analysis on the top 1000 cells based on the number of spatial clusters per cell (**Figure S1b**) (see **Methods**).

Next, we explored the genomic resolution achieved within each cell and we observed ∼40 reads within each 1-megabase bin across the genome for each individual cell (**Figure 1c**). Focusing on 20 randomly selected cells, we observed near-uniform coverage across each 1-megabase bin within the genome (**Figure S1c**).

Finally, we confirmed that scSPRITE accurately measures known genome structures. To do this, we merged the 1000 individual scSPRITE datasets (referred to as ensemble scSPRITE) and compared these to previously generated bulk SPRITE data in mouse ES cells (**Figure 1d**). We found that the scSPRITE maps are highly comparable to the bulk SPRITE data across various levels of resolution (Pearson correlation 0.97, Spearman correlation 0.94). Together, our results demonstrate that scSPRITE accurately measures genome organization and provides high resolution data on >1000 intact single nuclei.

### scSPRITE measures multiway interactions in single cells and generates high resolution single cell contact maps

Because each individual cell contains a single genome and contacts detected in multiple cells cannot be pooled together (as in bulk measurements), single cell genome structure methods need to maximize the number of contacts detected within each cell. This is the main challenge and limitation for all single cell genomic methods.

Currently, all existing single cell genome structure methods (e.g. scHiC) utilize proximity ligation and are therefore limited to measuring pairwise DNA contacts. In contrast, SPRITE captures multiway contacts among DNA molecules – rather than pairwise contacts – which dramatically increases the structural resolution that can be obtained for an individual cell (**Figure 2a**). Measuring multiple DNA molecules within one structure allows us to represent genomic features as structural blocks, as opposed to dots, which represent pairwise interactions in scHiC (**Figure 2a**). This is because the maximal number of interactions that can be captured increases quadratically with the size of a complex (**Figure S2a**). For example, if a crosslinked complex contains four DNA fragments, the maximum number of contacts that can be observed by pairwise methods is two, whereas the maximal number that can be identified with multiway contacts is six (**Figure 2a, Figure S2b**). Indeed, we observe an increase of >30-fold in contacts for each cell using scSPRITE (930,855,955/cell) compared to scHiC^16^ (375,470/cell) even though the number of sequencing reads per cell is <10-fold lower for scSPRITE relative to scHiC (**Figure 2b**).

**Figure 2:**
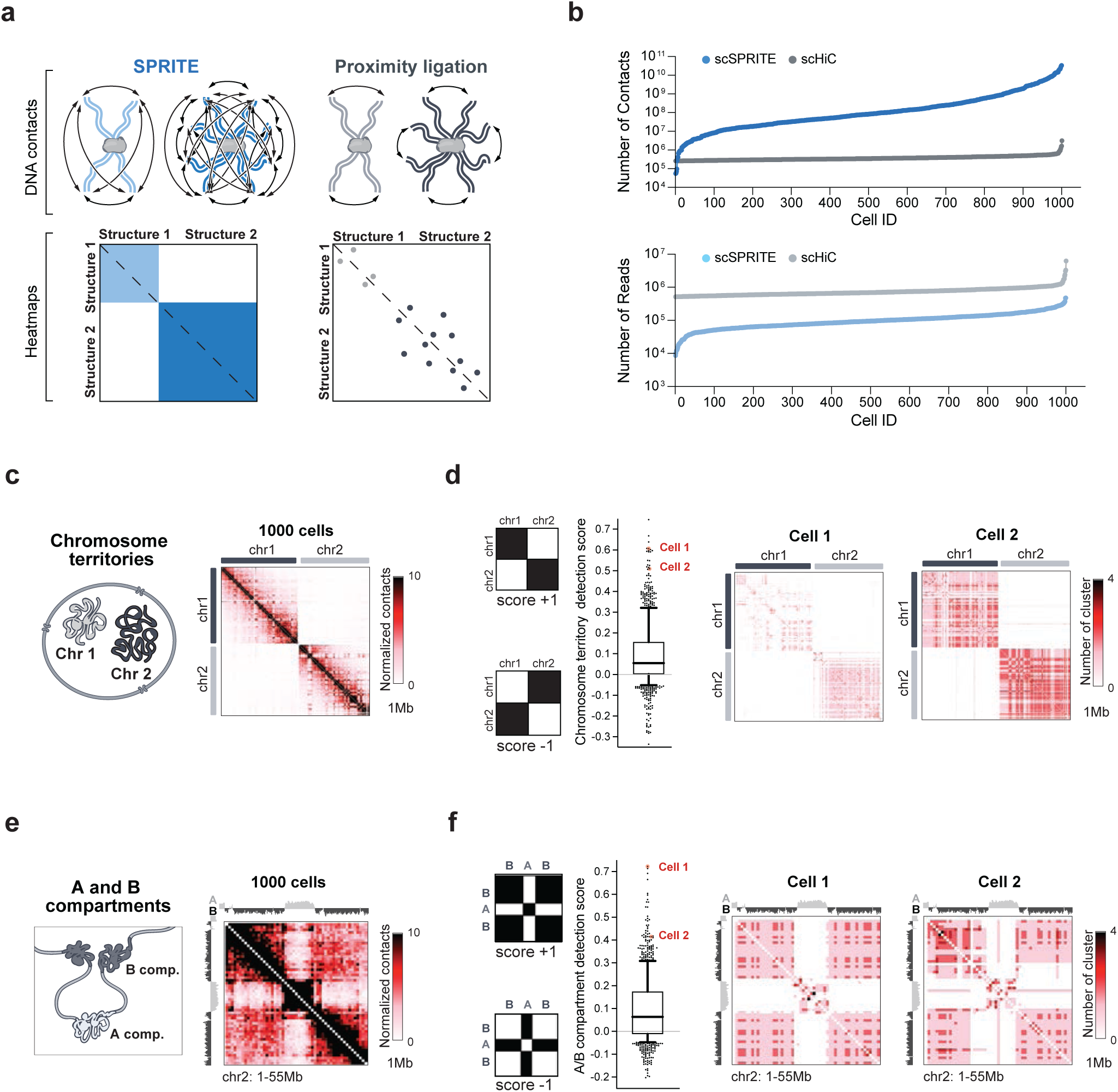
Capturing multiway interactions in single cells enables scSPRITE to accurately measure DNA interactions at different resolutions. **a**, Schematic illustration of differences between capturing pairwise interactions by multiway contacts (SPRITE-derived methods) and proximity ligation (HiC-derived methods) and examples of contacts maps. **b**, Comparison of number of contacts (top) and number of reads (bottom) obtained from scSPRITE (blue) and scHiC16 (grey) experiments rank ordered from smallest to largest. **c**, Representation of chromosome territory structure between chr1 and chr2: illustration (left) followed by ensemble scSPRITE heatmap (right); pairwise contact map at 1Mb resolution. **d**, Chromosome territory normalized detection scores for 1000 individual cells (left). To the left of the box plot, model representation of structures with max. score +1 and min. score -1. For the box plot, the whiskers represent the 10th and 90th percentiles and the central black line represents the median. In red, Cell1 and Cell2 (right) are demarcated. Cell1 and Cell2 are single cell examples of chromosome territories between chr1 and chr2. Contact map represents number of DNA clusters at 1Mb resolution. **e**, Representation of A/B compartment structure within 0-55Mb in chr2: illustration (left) followed by ensemble scSPRITE heatmap (right); pairwise contact map at 1Mb resolution. **f**, A/B compartments detection scores for 1000 individual cells (left). To the left of the box plot, model representation of structures with max score +1 and min. score -1. For the box plot, the whiskers represent the 10th and 90th percentiles and the central black line represents the median. In red, Cell1 and Cell2 (right) are demarcated. Cell1 and Cell2 are single cell examples of A/B compartments detected between 0-55Mb of chr2. Contact map represents number of DNA clusters at 1Mb resolution.

### scSPRITE detects chromosome territories and A/B compartments in single cells

To determine which DNA structures can be identified within single cells, we generated DNA contact maps from each of the 1000 individual cells. For every structure identified in the ensemble data, we computed a normalized detection score that reflects how well each single cell contact map resembles this structure compared to a randomized contact map. Briefly, for each structure we calculate an observed detection score which defines whether or not each pair of genomic bins within a structure were in contact. As a result, a single structure within a cell that contains all possible pairwise contacts would have a detection score=1 whereas a structure containing none of the expected pairwise contacts would have a detection score=-1. (see **Methods**). We normalized this observed scored to a distribution of scores generated by randomly permuting the locations of this structure.

We focused on genomic structures that were previously reported to occur within single cells – chromosome territories and A/B compartments^1^. Chromosome territories are structures containing high frequencies of intrachromosomal interactions with minimal interchromosomal interactions (**Figure 2c**). In our dataset, we looked at the contacts between chromosome 1 (chr1) and chr2, and detected clear separation of contacts into chromosome territories both in the ensemble data (**Figure 2c**) and in >75% of single cells (score>0, **Figure 2d, Figure S2c**). Genomes are further divided into A/B compartments, intrachromosomal structures defined by open (A) or closed (B) chromatin state (**Figure 2e**). Using scSPRITE, we detected the segregation of chromatin into A/B compartments in >65% of single cells (score>0, **Figure 2f, Figure S2d**). Together, our results demonstrate that scSPRITE can detect known genomic interactions in single cells.

### scSPRITE identifies inter-chromosomal DNA hubs organized around nuclear bodies

In addition to the known genomic structures discussed above, the nucleus is organized around various nuclear bodies that form higher-order inter-chromosomal contacts^7,8^. These interactions have not been previously explored within single cells because existing proximity-ligation methods do not detect multiple DNA molecules in one complex.

We explored whether scSPRITE can detect higher-order structures in single cells. First, we focused on interactions formed between centromere-proximal regions from different chromosomes. Centromeres and pericentromeres are long stretches of repetitive DNA that are essential for chromosome stability and segregation^25^, and have been shown to come into close proximity **(Figure 3a)** forming inter-chromosomal structures called chromocenters^25^. Because centromeres and peri-centromeres are not mapped in the genome, we focused our analysis on the first 10 Mb of each chromosome. In scSPRITE data we detect contacts between centromere-proximal regions of chr1 and chr11 both in the ensemble data (**Figure 3a**) and in >50% of single cells (score >0, **Figure 3b**). We also detected centromere-proximal interactions between other chromosome pairs (**Figure S3a**), showing that the formation of chromocenters can be analyzed in hundreds of single cells using scSPRITE.

**Figure 3:**
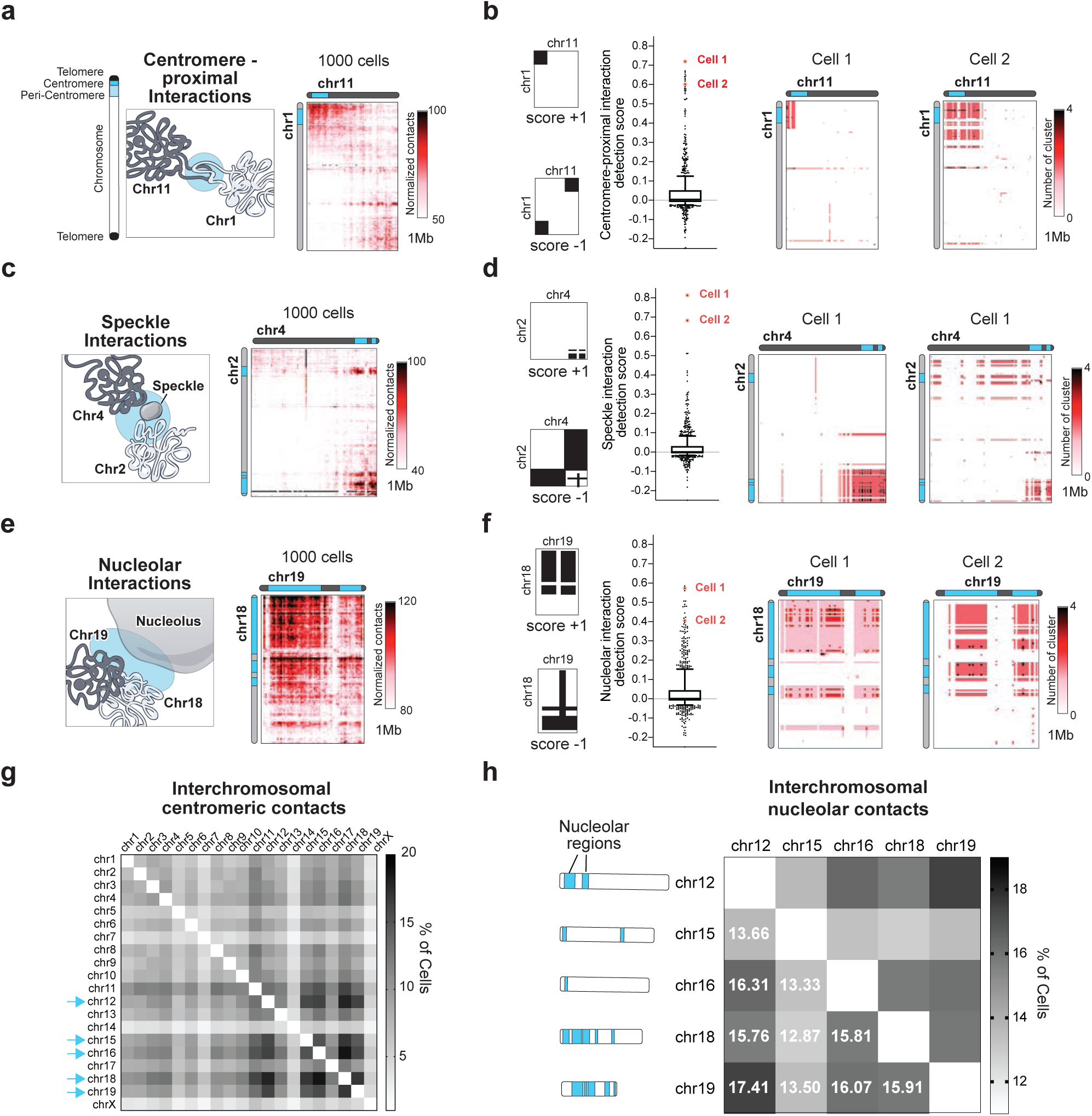
Higher-order structures are identified genome-wide in hundreds of single mESC by scSPRITE. **a**, Centromere-proximal interaction between chr1 and chr11: illustration (left) followed by ensemble scSPRITE heatmap (right); contact map at 1Mb resolution. **b**, Centromere-proximal region detection scores for 1000 individual cells (middle); the whiskers - the 10th and 90th percentiles, the central black line - median; single cell examples demarcated in red. Model representation of structures with max score +1 and min. score -1 (left). Single cell examples (right); contact map represents number of DNA clusters at 1Mb resolution. **c**, Speckle interaction between chr2 and chr4: illustration (left) followed by ensemble scSPRITE heatmap (right); contact map at 1Mb resolution. **d**, Speckle interaction detection scores for 1000 individual cells (middle); the whiskers - the 10th and 90th percentiles, the central black line - the median; single cell examples demarcated in red. Model representation of structures with max score +1 and min. score -1 (left). Single cell examples of speckle interaction detected between chr2 and chr4 (right); contact map represents number of DNA clusters at 1Mb resolution. **e**, Nucleolar interaction between chr18 and chr19: illustration (left) followed by ensemble scSPRITE heatmap (right); contact map at 1Mb resolution. **f**, Nucleolar interactions detection scores for 1000 cells (middle); the whiskers - the 10th and 90th percentiles, the central black line - the median; Single cell examples demarcated in red. Model representation of structures with max score +1 and min. score -1 (left). Single cell examples of nucleolar interactions detected between chr18 and chr19; contact map represents number of DNA clusters at 1Mb resolution. **g**, Heatmap representing the percent of cells containing interchromosomal centromeric contacts for each pair of chromosomes. Contacts normalized to number of reads per region. **h**, Heatmap representing the percent of cells containing inter-chromosomal nucleolar contacts for each pair of nucleolar associating chromosomes. Contacts normalized to number of reads per region.

We next focused on DNA organization around nuclear speckles. Nuclear speckles are structures enriched in pre-mRNA splicing factors that form higher-order interactions by bringing together gene-dense and highly transcribed Pol II regions from different chromosomes^11,26^. We focused on the previously reported inter-chromosomal interactions between precise regions of chromosomes 2 and 4 (**Figure 3c**) and found that >45% of cells contain this interaction (score>0, **Figure 3d**). We also detected speckle interactions between regions of chr2 and chr5 (**Figure S3b**), showing that speckle interactions can be detected at the single cell level by scSPRITE.

Finally, we focused on DNA interactions around the nucleolus, a nuclear body that is the location of ribosomal DNA (rDNA) transcription, ribosomal RNA processing, and ribosome biogenesis^12^. In mESCs, chr12, chr15, chr16, chr18, chr19 contain rDNA clusters and are known to form nucleolar interactions. Consistent with previous reports^7^, we identified strong nucleolar interactions between chr18 and chr19 in the ensemble scSPRITE dataset (**Figure 3e**). These interactions were also found in >50% of individual cells (score >0, **Figure 3f**). We also observed nucleolar contacts between another pair of rDNA-containing chromosomes, chr12 and chr19 (**Figure S3c**).

The results of these analyses demonstrate that scSPRITE can capture various higher-order contacts reflecting long-range interactions across multiple cells and involving structures of different sizes.

We note that centromere-proximal and nucleolar contacts were not detectable even in the ensemble scHiC data^16^ (**Figure S3d**). Although the ensemble scHiC was able to identify speckle interactions, the single-cell interaction maps lacked information on these structures.

### scSPRITE reveals cell-to-cell heterogeneity of higher-order structures in mESCs

Single-cell measurements capture dynamics in genome structure across a population of cells. To explore this heterogeneity, we looked at the frequency of centromere-proximal regions (**Figure 3g**) and nucleolar interactions (**Figure 3h**). Each centromere-proximal region formed a pair with another chromosome in ∼8% of cells, with the exception of chr7, chr14, and chrX, which interacted with another chromosome’s centromere-proximal region in <5% of cells (**Figure S3e**). Interestingly, we observed that centromere-proximal regions of mESC chromosomes that contained nucleolar organizing regions (NORs) tended to interact with each other more frequently (>12% of cells) than with other chromosomes (**Figure 3g**). This type of nuclear organization (where NORs cluster their centromere and peri-centromere regions around nucleoli to form chromocenters) has been previously described by microscopy^27,28^.

Next, we looked at nucleolar contacts and measured the frequency of interactions between different pairs of NOR regions in each chromosome. We detected the most NOR region interactions between chr19 and chr12 (17.41%), and the fewest interactions between chr15 and chr18 (12.87%) (**Figure 3h**). Overall, each NOR-containing chromosome formed a pair with another NOR-containing chromosome in at least ∼15% of cells, with the exception of chr15, which interacted the least frequently (∼12% of cells) (**Figure S3f**).

### scSPRITE demonstrates that TADs are heterogeneous units of mESC genome organization

Topologically associating domains (TADs) are intrachromosomal structures in which contiguous regions of the genome have been shown to interact more with themselves than with surrounding regions^2,3,29^, however their global existence in single cells has been debated ^13,15,16,20^. It is unclear whether this reflects technical limitation of current single cell methods (e.g. low-resolution structures) or if these DNA structures are not present in individual genomes. Because scSPRITE generates higher resolution structures within individual cells, we asked whether TADs can be detected in single cells with our method.

First, we focused our analysis on a region of chr4 where we observed strong evidence for TADs in the ensemble scSPRITE dataset (**Figure 4a**), and we detected similar TAD-like structures in >75% of single cells (score>0, **Figure 4b**), suggesting that this TAD is present in the majority of individual cells. Next, we asked whether we could detect more TAD-like structures across the genomes of single cells. We looked at a handful of strong TADs from our ensemble dataset and observed their clear presence in single cells (**Figure S4a**). We scored all of the identified TADs (from our ensemble dataset) in each of 1000 individual cells and averaged the score per cell; we found that in the majority of cells we can identify structures similar to TADs (>90% cells with score>0, **Figure S4b**). These results demonstrate that TAD-like structures can be detected within individual cells.

**Figure 4:**
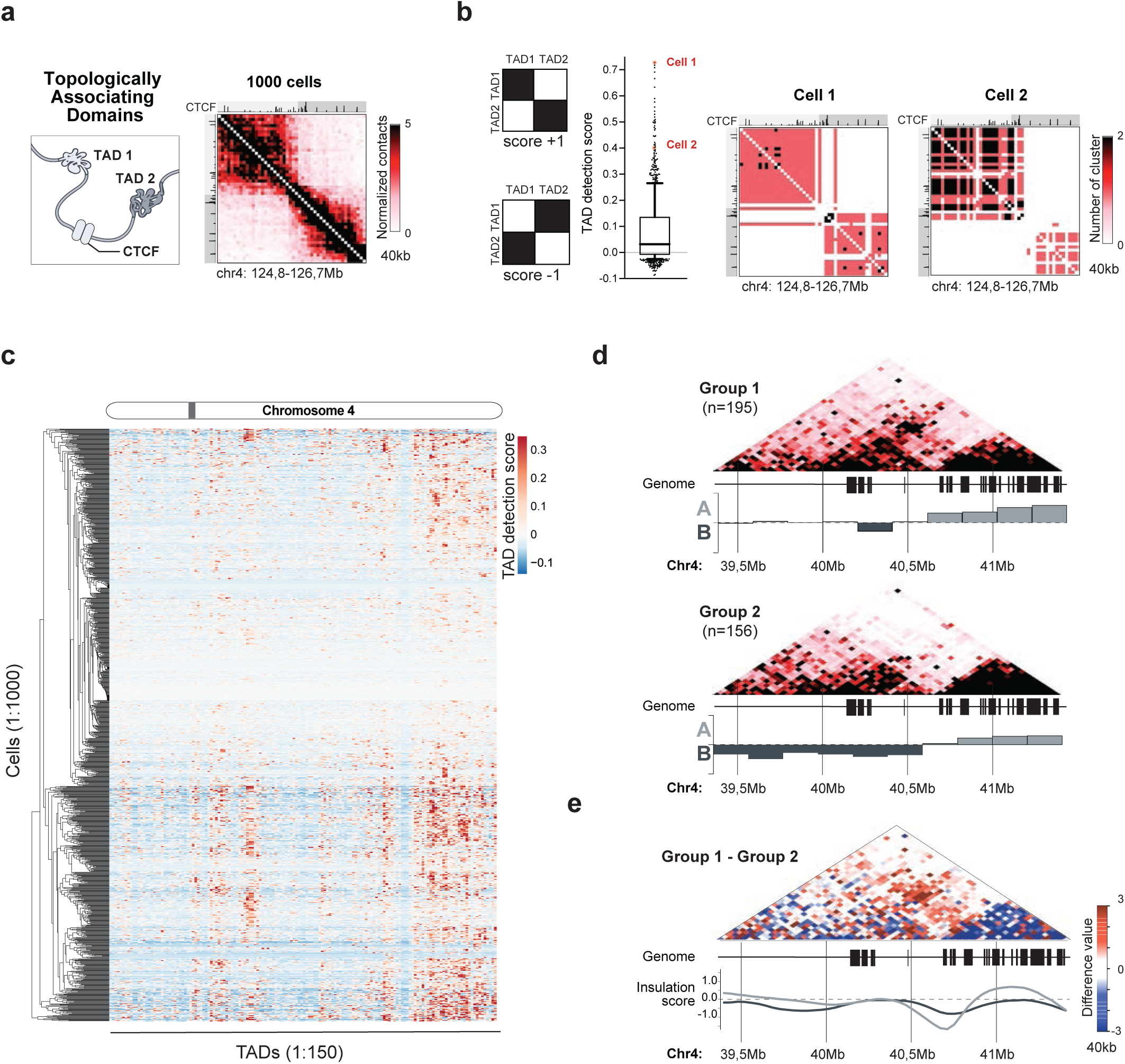
TADs are heterogeneous units present in the genomes of individual mESCs. **a**, TAD structure between 124.8-126.7Mb of chr4: illustration (left) followed by ensemble scSPRITE heatmap (right); pairwise contact map at 40kb resolution. **b**, TAD detection scores for 1000 cells (middle); the whiskers - the 10th and 90th percentiles, the central black line - the median; single cell examples demarcated in red. Model representation of structures with max score +1 and min. score -1 (left). Single cell examples (right); contact map represents number of DNA clusters at 40kb resolution. **c**, TAD detection scores across 1000 cells: columns represent the strength of TAD detection scores for all TADs detected across chr4 in ensemble scSPRITE, rows represent each single cell from 1000 analyzed cells clustered based on score similarity pattern; grey bar - variable region described in the text and zoomed in Figure S4e. **d**, Ensemble heatmaps across 39.4-41.4Mb region of chr4 representing cells containing (Group I, top) or lacking (Group I, bottom) the contact emerging over the boundary of A/B compartment. For each set, the eigenvector was calculated to represent A/B compartments strength. **e**, Difference contact map across 39.4-41.4Mb of chr4 made by subtracting the normalized contacts in Group II from Group I (Figure 4d). Insulation scores for cells in Group I (dark grey) and Group II (light grey) are plotted.

Next, we sought to understand whether TADs are stable units of genome organization or whether they are heterogenous structures among cells. We looked at the distribution of TAD detection scores per individual TAD by averaging scores from 1000 single cells and observed that >65% of TADs scored below 0 (**Figure S4c**) suggesting that they were not present in many individual cells.

To determine if the differences in scores depend on genomic location and/or specific cell, we scored each of the 1000 cells individually for all detected TADs (**Figure 4c, Figure S4d)**. We observed variability in TAD detection scores among cells, but we also found genomic regions spanning several TADs with well-defined TADs (score >0) and with no TADs (score <0). This observation suggests that some genomic regions have more clearly defined TADs in a single cell than others.

We examined one of the variable regions (**Figure 4c, Figure S4e**) in more detail and found a weak TAD emerging in the ensemble scSPRITE dataset over the A/B compartment boundary (**Figure S4f**). We analyzed cells with coverage over the region of interest and identified two groups of cells with different TAD-like structures and shifted A/B compartment boundaries (**Figure 4d**). We observed a structure emerging across boundaries only in one group of cells and the TAD insulation scores for two groups were shifted (**Figure 4e**). These distinct TAD structural states cannot be simply explained by differences in cell cycle (**Figure S4g**) or by other major structural changes between two groups of cells (**Figure S4h**).

### scSPRITE detects structural heterogeneity in promoter-enhancer interactions

We next asked if scSPRITE could detect changes in TADs that reflect biologically significant long-range DNA contacts, such as the interactions between promoters and super enhancers (SE). SE are large domains enriched in histone 3 lysine 27 acetylation (H3K27ac) that are thought to modulate gene expression by forming loops with promoters^6^. Bulk genome-wide studies have shown that SE can form long- and short-range interactions with the same promoter^30–32^, but it remains unclear whether these interactions occur simultaneously in the same cell. We focused on the Nanog locus, a key pluripotency factor in ES cells whose promoter interacts with multiple enhancers over broad range of distances (up to 300 kb)^30,33^ (**Figure 5a**). We selected cells with coverage over the region of interest, split cells into two groups according to whether contact with the SE 300 kb upstream of the Nanog locus (−300 kb SE) was detected or not (**Figure S5a**), and observed changes in TAD-like structures. Surprisingly, we noticed that in the group with long-range interactions detected, short-range interactions were weaker (**Figure 5b**). We confirmed that these observations were not caused by differences in cell cycle phase (**Figure S5b**). This suggests that when the long-range Nanog-SE contact is present, cells are less likely to form short-range contacts (and vice versa).

**Figure 5:**
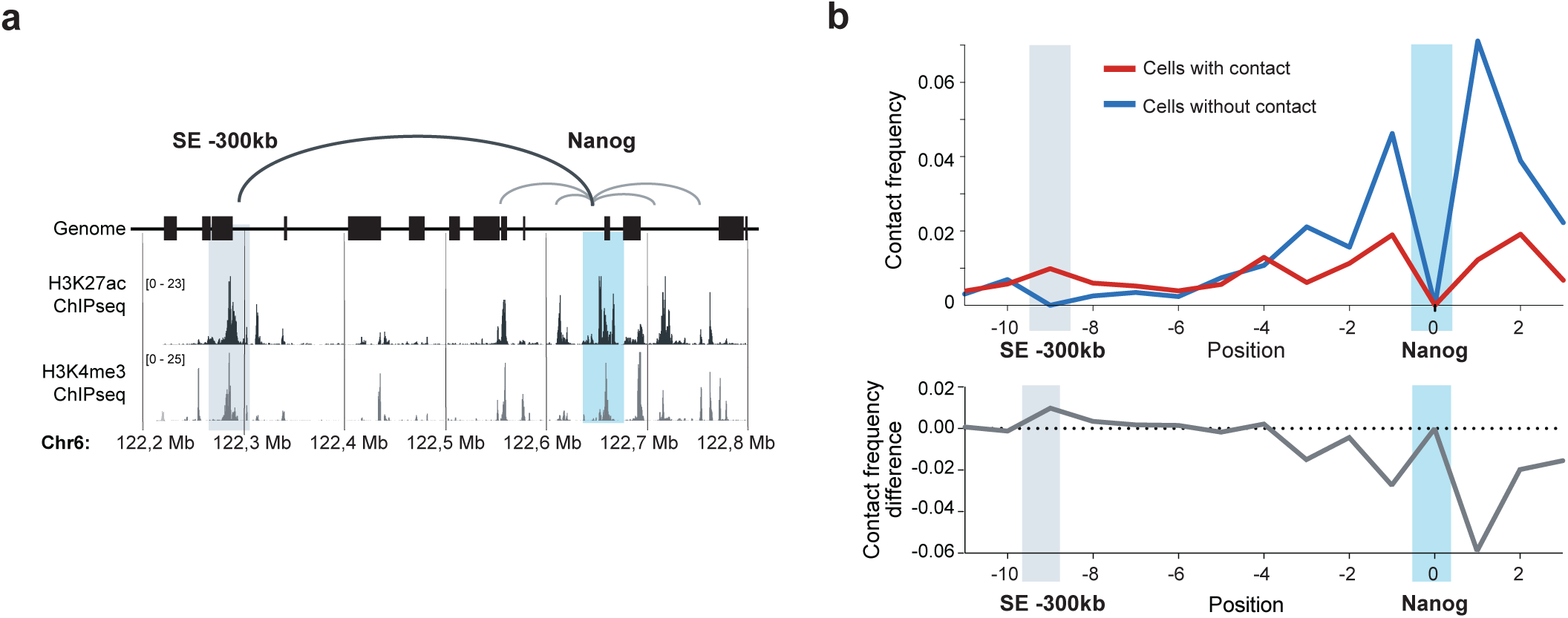
The Nanog locus in mESCs represents heterogeneous structural state forming short- or long-range interactions with its SEs. **a**, Representation of Nanog locus and its DNA interactions with SE: 122.2-122.8 Mb region in chr6 with corresponding ChIPseq tracks for H3K27ac and H3K4me3; Nanog-SE interactions (grey lines). **b**, Normalized contact frequency plot between Nanog locus and 122.2-122.8 Mb surrounding region in chr6; Shown cells containing (red) or lacking (blue) the contact between the Nanog locus and SE -300 kb (top); Below, difference in normalized contact frequency plotted (bottom).

Next, we looked at the previously described^32^ long-range interaction between the Tbx3 gene and an enhancer region near the Lhx5 gene, which are 760 kb apart. We analysed cells with coverage over the region of interest, split cells into two groups (one with and the other without interactions), and observed differences in TAD-like structures between the two groups (**Figure S5c, S5d**), suggesting that the Tbx3-Lhx5 contact is also heterogenous in mESCs.

## DISCUSSION

Here we describe scSPRITE, a method to generate high-resolution genome-wide maps of genome organization in thousands of single cells simultaneously. Our results reveal several novel insights into genome organization and the heterogeneity of genome structure.

(i) To date, most genome-wide methods have focused on short-range and intrachromosomal interactions; using scSPRITE we identified long-range higher-order interactions of both active (nuclear speckle) and inactive (centromeres and nucleolar contacts) chromatin regions in hundreds of single cells. These results open a new path to dissect function and heterogeneity of DNA contacts around nuclear bodies in individual cells.

(ii) Although results from bulk methods have led to the proposal that TADs form stable, functional units of chromosome organization^34–36^, our single cell data demonstrates that TADs differ within individual cells. Moreover, our data suggests that some genomic regions have more clearly defined TADs in a single cell than others. These global observations are consistent with recent microscopy data that studied specific loci ^13,14,20^.

(iii) Although biologically relevant DNA interactions like promoter-enhancer contacts were previously thought to occur within TAD boundaries ^29,34,35^, our results demonstrate that the action of regulatory sequences may not be restricted within one structural unit. For example, we showed that the Nanog locus forms long-range interactions with its -300 kb SE across its TAD boundary or short-range interactions with proximal SE within TAD region.

Because scSPRITE is simple to perform – it does not require specialized equipment, techniques, or training – we expect that it will greatly expand the availability of single cell genome structure measurements to any laboratory with minimal molecular biology background and facilitate single cell studies across a broad spectrum of cell-types. Importantly, scSPRITE can be scaled to work with as few as hundreds or up to thousands of cells simultaneously. It can be adapted to work with different cell types or homogenized tissues that are composed of mixed cell populations, without a need for laborious and ineffective cell purification to homogeneity.

Currently, one of the main challenges with studying complex tissues (e.g. brain) or diseases (e.g. tumors) is the heterogeneity of their cellular composition. The application of scSPRITE to such mixed cell populations will enable studies of intrinsically heterogeneous systems and will provide a global and accurate view of their 3D genome organization. Accordingly, we expect that scSPRITE will provide the field with a path toward understanding the relationship between 3D genome organization and genome function within single cells.

## METHODS

### Cell types and culture conditions

We developed scSPRITE using mouse and human cells, focusing primarily on mouse embryonic stem cells (mESCs) because their genome structure has been extensively studied^3,15^. Because mESCs display both phenotypic and functional heterogeneity^17,23,24^, they provide a good model system for studying cell-to-cell variability with single-cell measurements.

Mouse embryonic stem (ES) cell lines were cultured in serum-free 2i/LIF medium and maintained at an exponential growth phase as previously described^37^. SPRITE maps were generated in female bsps ES cell line (provided by K. Plath, UCLA), a female ES cell line derived from the V6.5 ES cell line, which expresses Xist from the endogenous locus under the transcriptional control of a tet-inducible promoter and the Tet transactivator (M2rtTA) from the Rosa26 locus.

HEK293T, a female human embryonic kidney cell line transformed with the SV40 large T antigen was obtained from ATCC (#CRL-1573) and cultured in complete media consisting of DMEM (#11965092, GIBCO, Life Technologies; Carlsbad, CA) supplemented with 10% FBS (Seradigm Premium Grade HI FBS, VWR), 1X penicillin-streptomycin (GIBCO, Life Technologies), 1X MEM non-essential amino acids (GIBCO, Life Technologies), 1 mM sodium pyruvate (GIBCO, Life Technologies) and maintained at 37°C under 5% CO2. For maintenance, 800,000 cells were seeded into 10 mL of complete media every 3-4 d in 10-cm plates.

### scSPRITE protocol

#### Cell crosslinking

Media from adherent mESCs (bsps) was removed and washed once with 1X PBS. Cells on the 10 cm plates were then trypsinized using 2 mL of 0.025% Trypsin-EDTA (prewarmed to 37°C). Plates were incubated at 37°C for 5 min and the trypsinized cells were mixed by pipetting to break up any clumps. We added 8 mL of pre-heated wash solution (DMEM/F12 + BSA, prewarmed to 37°C) to the plate to inactivate trypsin before transferring the cells to a conical tube. Cells were centrifuged at 330 *g* for 3 min, and the supernatant was discarded. Cells were washed once with 1X PBS at a ratio of 4 mL of PBS per 1 × 10^7^ cells and centrifuged again at 330 *g* for 3 min. After the wash, 4 mL of 2 mM disuccinimidyl glutarate (DSG,Life Technologies, #20593, Carlsbad, CA) prepared in 1X PBS was added per 1 × 10^7^ cells to the conical tube and the solution was mixed thoroughly by pipetting to remove clumps. The cells in DSG solution were gently shaken for 45 min at room temperature. Following incubation with DSG, 200 µL of 2.5 M glycine were added per 1 mL of DSG solution previously added to quench the reaction, and the tube was gently shaking for 5 min at room temperature. Cells were then centrifuged at 1000 *g* for 4 min and the supernatant was discarded. Cells were washed with 1X PBS at a ratio of 4 mL of PBS per 10 M cells and centrifuged again at 1000 *g* for 4 min. After the wash, 4 mL of 1% formaldehyde (16% (w/v) formaldehyde ampules, Life Technologies, #28908, Carlsbad, CA) prepared in pre-warmed (37°C) 1X PBS was added per 1 × 10^7^ cells to the conical tube and the solution was mixed thoroughly by pipetting to remove clumps. The cells in formaldehyde solution was then gently shaking for 10 min at room temperature. Following incubation with formaldehyde, 200 µL of 2.5 M glycine was added per 1 mL of formaldehyde solution previously added to quench the reaction, and the tube was gently shaking for 5 min at room temperature. Cells were then centrifuged at 1000 *g* for 4 min and supernatant was removed. Cells were twice washed with cold 1X PBS + 0.5% BSA (w/v) solution, and centrifugation was done at 4°C at 1000 *g* for 4 min. Following the washes, enough cold 1X PBS + 0.5% BSA solution was added to get a cell concentration of 5 × 10^6^ cells/mL. Crosslinked cells were then aliquoted in new 1.5 mL low-bind Eppendorf tubes, centrifuged (2000 *g* for 5 min) to remove the supernatant, and flash-frozen in liquid nitrogen. Cells were kept at - 80°C until used for analyses.

#### Cell lysis and nuclei preparation

Cells crosslinked previously were thawed from -80°C and were kept on ice during the cell lysis procedures. Initially, 1.4 mL of lysis buffer #1 (50 mM HEPES pH 7.4, 1 mM EDTA pH 8.0, 1 mM EgTA pH 8.0, 140 mM NaCl, 0.25% TritonX-100, 0.5% IGEPAL CA-630, 10% glycerol, 1X proteinase inhibitor cocktail (PIC)) was added per 1 × 10^7^ cells. The cell solution was mixed thoroughly before incubating on ice for 10 min. Cells were pelleted afterwards at 900 *g* for 8 min at 4°C and the supernatant was removed. Following, 1.4 mL of lysis buffer #2 (10 mM Tris-HCl pH 8, 1.5 mM EDTA, 1.5 mM EgTA, 200mM NaCl, 1X PIC) was added per 1 × 10^7^ cells. Again, the cell solution was mixed thoroughly before incubating on ice for 10 min. Cells were pelleted afterwards at 900 *g* for 9 min at 4°C and supernatant was removed. Afterwards, the cells were washed in 800 µL of 1.2X CutSmart solution (from 10X CutSmart stock (NEB, #B7204S, Ispwich, MA)) and pelleted at 900 *g* for 2 min. Supernatant was removed and a fresh 400 µL of 1.2X CutSmart solution was added carefully to not resuspend the pellet. Then, 6 µL of 20% SDS was added to the tube, and the cells were thoroughly resuspended. The cell solution was mixing on an Eppendorf ThermoMixer C at 1200 rpm for 60 min at 37°C to isolate nuclei. Next, 40 µL of 20% Triton X-100 was added to the same tube to quench the reaction, and the solution was left mixing on the same instrument at 1200 rpm for 60 min at 37°C. Lastly, 30 µL of 5000 U/mL HpyCH4V (NEB, #R0620L, Ispwich, MA) was added to the same tube to allow for DNA to be digested in-nuclei. In-nuclei digestion was performed for 4 h at 37°C while shaking at 1200 rpm. Following digestion, nuclei were pelleted at 900 *g* for 2 min, supernatant was removed, and nuclei washed three times with 1X PBS, 1 mM EDTA, 1 mM EgTA, and 0.1% Triton X-100 solution at 900 *g* for 2 min. Following the washes, the nuclei concentration was assessed by loading 6 µL of the solution into a disposable hemocytometer (4-Chip Disposable Hemocytometer, Bulldog Bio, #DHC-N420, Portsmouth, NH). After determining nuclei concentration, 5 × 10^5^ nuclei were transferred by pipetting into a new 1.5 mL low-bind Eppendorf tube. In this new tube, 25 µL of dA-tail reaction buffer and 10 µL of Klenow Fragment were added to the nuclei (both reagents were part of NEBNext dA-Tailing Module (NEB, #E6053L, Ispwich, MA)). The tube was filled to 250 µL using nuclease-free H_2_O and dA-tailing was performed in-nuclei at 37°C for 90 min while shaking at 1200 rpm. The reaction was then stopped with the addition of 200 µL of 1X PBS, 50 mM EDTA, 50 mM EgTA, and 0.1% Triton X-100. The nuclei pellet was spun down at 900 *g* for 2 min and washed twice using 400 µL of 1X PBS, 1 mM EDTA, 1 mM EgTA, and 0.1% Triton X-100 solution. Following the washes, the nuclei were resuspended in fresh 1X PBS, 1 mM EDTA, 1 mM EgTA, and 0.1% Triton X-100 solution and nuclei concentration was determined again using the hemocytometer as described previously.

#### In-nucleus combinatorial barcoding

To uniquely identify DNA sequences originating from the same cell, combinatorial barcoding was performed in-nuclei. Three rounds of combinatorial barcoding were performed in the following order: “DNA phosphate modified” (DPM), “odd” tagging, and “even” tagging (these tags are described in the original SPRITE paper^7^. The resulting tags were pre-loaded onto a 96-well plate, with each well containing 2.4 µL of a uniquely barcoded tag at a concentration of 45 µM. Nuclei previously dA-tailed were washed twice in a solution of 1X PBS, 0.1% Triton X-100, and 0.3% BSA (w/v), and nuclei concentration was reassessed using a hemocytometer, as described previously. To perform in-nuclei barcoding, 2 × 10^5^ nuclei were withdrawn and transferred into a new 1.5 mL low-bind Eppendorf tube, and filled to 1125 µL using a solution of 1X PBS, 0.1% Triton X-100, and 0.3% BSA (w/v). The nuclei solution was well-mixed before loading 11.2 µL of nuclei solution into each well of a 96-well plate. Each well was then supplemented with 6.4 µL of ligation mix (220 µL of 2X Instant Sticky Master Mix (NEB, #M0370, Ispwich, MA), 352 µL 5X Quick Ligase Buffer (NEB, #B6058S, Ispwich, MA), and 132 µL 1,2-Propanediol (Sigma, #398039, St. Louis, MO)). The 96-well plate was sealed after loading ligation mix and was mixed on an Eppendorf ThermoMixer C at 20°C. The reaction was performed for 3 h while mixing at 1600 rpm for 30 s every 5 min. After performing in-nuclei DNA ligation, 20 µL of 1X PBS, 50 mM EDTA, 50 mM EgTA, and 0.1% Triton X-100 solution was added to each well and incubated for 10 min at 20°C to stop the ligation reaction. Next, a solution of 80 µL of 1X PBS, 50 mM EDTA, 50 mM EgTA, and 0.1% Triton X-100 (w/v) was added to each well, and all the contents of the well plate were pooled together into a new 15 mL conical tube. The 96-well plate was washed once with a solution of 100 µL of 1X PBS, 50 mM EDTA, 50 mM EgTA, and 0.1% Triton X-100 and pooled together into the same conical tube. Nuclei were pelleted at 800 *g* for 10 min, and all but 1 mL of supernatant was removed from the tube. The nuclei were resuspended before transferring to a new 1.5 mL non-low bind Eppendorf tube. In the new Eppendorf tube, nuclei were washed twice with a solution of 500 µL of 1X PBS, 0.1% Triton X-100, and 0.3% BSA (w/v) at 900 *g* for 2 min. This in-nuclei ligation process was repeated until the “DPM”, “odd”, and “even” tags were added sequentially.

#### Sonication

Once the in-nuclei barcoding process was completed, nuclei were filtered through a 10 µm mesh filter (PluriStrainer, #43-10010-50, Spring Valley, CA) into a new 1.5 mL non-low bind Eppendorf tube to ensure we only isolated single cells (**Figure S1a**). Filtered nuclei were then pelleted at 900*g* for 2 min and supernatant was removed. Nuclei were resuspended and washed twice in lysis buffer #3 (1.5 mM EDTA, 1.5 mM EgTA, 100 mM NaCl, 0.1% sodium deoxycholate, 0.5% sodium lauroyl sarcosinate) at 900*g* for 2 min. The nuclei concentration was determined again using a hemocytometer. From there, 1500 nuclei were removed and placed into a Covaris microtube-15 and filled to 15 µL using lysis buffer #3. The Covaris tube was placed in the Covaris M220 Focused-ultrasonicator (Covaris, Woburn, MA) and sonication was performed for 2 min under specific settings (water temperature 6°C, incident power 30W, duty cycle 3.3) to release DNA complexes from nuclei. The tube was then removed from the instrument and set on ice.

#### NHS (*N-*hydroxysuccinimide) beads coupling

After sonication, NHS-Activated Magnetic Beads (Life Technologies, #88826, Carlsbad, CA) were activated for coupling. First, 600 µL of NHS-beads were withdrawn and placed in a 1.5 mL low-bind Eppendorf tube. The tube was placed on a DynaMag-2 magnet, and the supernatant was removed. The beads were washed once with 600 µL of ice-cold 1M HCl, and the supernatant was removed again and replaced with 600 µL of ice-cold 1X PBS. After removing 1X PBS, the beads were resuspended in 500 µL of 1X PBS + 0.1% SDS. Additionally, 85 µL of 1X PBS + 0.1% SDS was added to the previously sonicated nuclei solution, mixed, and added to the bead solution. The complexes were then coupled to NHS-beads on an Eppendorf ThermoCycler C overnight at 4°C while shaking at 1200 rpm. After coupling, the flowthrough was removed and 600 µL of 1M Tris-HCl pH 7.5, 0.5 mM EDTA, 0.5 mM EgTA, and 0.1% Triton X-100 was added to the beads to quench the remaining NHS groups; this was done at 4°C at 1200 rpm for 60 min. Once the beads were quenched, the flowthrough was removed, and the beads were washed twice in cold RLT2+ buffer (0.2% Sodium lauryl sarcosinate, 1 mM EDTA, 1 mM EgTA, 10 mM Tris-HCl pH 7.5, 0.1% Triton X-100, 0.1% NP-40, filled to the final volume with RLT (Qiagen, #79216, Valencia, CA)). This was followed by three washes in M2 buffer (50 mM NaCl, 20 mM Tris-HCl pH 7.5, 0.2% Triton X-100, 0.2% NP-40, 0.2% sodium deoxycholate). The beads were then resuspended in a mix of M2 buffer and H_2_O (58% M2, 42% H_2_O) to attain a total volume of 1125 µL of M2 buffer, H_2_O, and beads.

#### Spatial barcoding/complex-specific barcoding

The complexes on beads underwent split-pool barcoding as described previously (Quinodoz et al. *Cell* 2018) to ligate three additional tags (odd, even, and Y-even) to the previously applied cell-specific tags. The bead solution was well-mixed before loading 11.2 µL of bead solution into each well of a 96-well plate, with each well containing 2.4 µL of a uniquely barcoded tag at a concentration of 4.5 µM. Each well was then supplemented with 6.4 µL of ligation mix (220 µL of 2X Instant Sticky Master Mix (NEB, #M0370, Ispwich, MA), 352 µL 5X Quick Ligase Buffer (NEB, #B6058S, Ispwich, MA), and 132 µL 1,2-Propanediol (Sigma, #398039, St. Louis, MO)). The 96-well plate was sealed after loading ligation mix and was mixed on an Eppendorf ThermoMixer C at 20°C. The reaction was performed for 60 min with mixing at 1600 rpm for 30s every 5 min. Afterwards, the reaction was stopped by adding 60 µL of RLT2+ buffer to each well before pooling the solutions of each well into a 25 mL reservoir. Each well was then rinsed once with 100 µL of RLT2+ buffer to remove residual beads and pooled into the same 25 mL reservoir. The solution was then transferred to a 15 mL conical tube, which was then placed on a magnet to remove most of the RLT2+ buffer from the beads. With about 2 mL of RLT2+ buffer remaining, the beads were resuspended and transferred to a low-bind 1.5 mL Eppendorf tube, which was placed on a DynaMag-2 magnet to remove the remaining RLT2+ buffer. The beads were washed three times with 600 µL of M2 buffer. This process of split-pool barcoding on beads was repeated until the three additional tags were added. After the last round of split-pool barcoding was completed, the beads were resuspended in 600 µL of MyK buffer (20 mM Tris-HCl pH 8.0, 0.2% SDS, 100 mM NaCl, 10 mM EDTA, 10 mM EgTA, 0.5% Triton X-100) following the washes.

#### Library Preparation

To ensure we capture all information coming from single cells, we need to sequence all DNA molecules that were bound to the beads. The bead solution was split equally into 10 low-bind Eppendorf 1.5 mL tubes, with each tube containing 60 µL of beads in MyK buffer. Next, an additional 32 µL of MyK buffer and 8 µL of Proteinase K (NEB, #P8107S, Ispwich, MA) were added to each tube. All 10 tubes were placed on an Eppendorf ThermoCycler C, and reverse crosslinking proceeded overnight at 60°C while shaking at 1200 rpm. Next, the tubes were placed on a DynaMag-2 magnet, and the MyK and Proteinase K solution were transferred to 10 new low-bind Eppendorf tubes. The beads from each of the tubes were washed once with 20 µL of H2O and then transferred to the same tube containing each respective MyK and Proteinase K solution. DNA from each of the tubes were purified using the Clean-and-Concentrator-5 columns (Zymo, #D4004, Irvine, CA) using 5X binding buffer to increase yield. Purified DNA from each column was eluted in 10 new Eppendorf 1.5mL tubes using 12 µL of water. Each of the tubes were filled to 30 µL using 15 µL Q5 Hot Start High-Fidelity 2X Master Mix (NEB, #M0493S, Ispwich, MA), 1.5 µL of 20X Evagreen (Biotium, #31000-T, Fremont, CA), 1.2 µL of 25 µM indexed Illumina primers, and 0.3 µL of H_2_O. Real-time PCR amplification proceeded for 14 cycles, which was when the libraries entered exponential amplification but had not plateaued. Following amplification, each of the libraries was diluted 4-fold prior to running on a 1% Agarose E-gel (Life Technologies, #G402001, Carlsbad, CA) with a E-Gel 1-Kb Plus DNA Ladder (Life Technologies, #10488090, Carlsbad, CA) as a reference. After the run, the gel was cut between 300 and 1000 bp marks to remove primer dimers, small non-specific amplicons, and long DNA amplicons. Libraries from the gel were purified using a Gel Purification Kit (Zymo, #D4002, Irvine, CA) as described by the manufacturer, and 20 µL of H_2_O was used to elute libraries off the column. To estimate the number of unique molecules in our libraries, the molarity of our libraries was determined using the concentration of our library from Qubit 3.0 Fluorometer (using the Qubit dsDNA high-sensitivity assay kit) and the average library size (bp) using an Agilent Tapestation 2200 (using the Agilent high-sensitivity D1000 ScreenTape and reagents). This in addition to estimated losses during library cleanup allowed us to estimate the number of unique molecules in our libraries. The libraries were sequenced with a read depth of 2.4X to ensure that most molecules were uniquely sequenced.

#### scSPRITE Data Generation & Sequencing Analysis

scSPRITE data was generated using Illumina paired-end sequencing on the Novoseq through Novogene Corporation. Reads were sequenced with at least 120 bp in Read 1 for genomic DNA information and the DPM tag and 95 bp in Read 2 to read the other five remaining tags (odd – even - odd - even - Y-even).

#### Sequencing pipeline analysis

scSPRITE barcodes were identified by combining the DNA tag sequence from the beginning of Read 1 and the remaining five barcode tags from Read 2. The tags were identified from a table of known tag sequences as previously described^7^, with Odd and Even tags allowing up to two mismatches and DPM and Y-even tags allowing zero mismatches. Any reads that lacked the full six barcode sequence (DPM - odd - even - odd - even - Y-even) in the expected order was discarded from further analysis. Before alignment, Read 1 was trimmed to a length of 100 bp.

#### Alignment and filtering of reads

The trimmed reads containing the full six barcode sequence were mapped to pre-indexed mm9 reference genome using STAR 2.6.1 using the following parameters: --outFilterMultimapNmax 50 --outFilterScoreMinOverLread 0.30 --outFilterMatchNminOverLread 0.30 -- outFilterIntronMotifs None --alignIntronMax 50000 --alignMatesGapMa× 1000 --genomeLoad NoSharedMemory --outReadsUnmapped Fastx --alignIntronMin 80 --alignSJDBoverhangMin 5 --sjdbOverhang 100 --limitOutSJcollapsed 10000000 --limitIObufferSize=300000000. SAMtools 1.9 was applied to filter mapped reads, and only uniquely mapped reads (-q 255) were kept. Alignments that had overlapped a masked region as denoted by Repeatmasker (UCSC, milliDiv < 140) were removed using bedtools (version 2.25.0). Finally, reads that were aligned to a non-unique region of the genome were removed by excluding alignments that mapped to regions by the ComputeGenomeMask program (read length = 35 nt). After these filtration steps, all BAM files that corresponded to the same sample but contained different Illumina primers at sequencing were pooled together before cluster identification.

#### Cluster barcode and cell barcode identification

To identify SPRITE clusters, all reads that contained the same six barcode sequences were grouped together into a single cluster. All reads containing the same six barcode sequence that started at the same genomic position were removed to remove possible PCR duplicates. Once identified, a SPRITE cluster file was generated where each line contained the cluster barcode name and corresponding genomic alignments. Once the cluster barcodes were identified, the cell barcodes were identified by grouping clusters together that contained the same DPM, first Odd, and first Even barcode sequences. This grouping can create on the order of hundreds of thousands cell barcode files, but the majority of these files contain fewer than 10 clusters. As a result, only the largest 4000 cell barcode files based on file size were selected for downstream filtration, and the remaining cell barcode files were removed from the directory.

### Data analysis

#### Selecting single cells for analysis

Once the largest 4000 cell barcode files were identified, these files underwent additional *in silico* filtration to select the appropriate cells for analysis. The files were rank-ordered based on the number of clusters. The 1500 cell barcode files with the largest number of clusters from the initial 4000 files were selected, consistent with initial number of cells used for the scSPRITE experiment. To ensure we selected single cells for downstream analysis, the top 3.4% percent of cells, as determined from the results of the human-mouse mixing experiment, were removed. From the remaining files, to ensure that the analysis containing the highest coverage per cell were analyzed, we then selected the top 1000 cell barcode files containing the most number of clusters per cell for downstream single-cell analysis.

#### Comparison of scSPRITE and SPRITE data

To compare with the SPRITE dataset, contact maps from scSPRITE were normalized using Hi-Corrector^38^ to generate maps at different genomic resolutions.

#### Insulation scores and A / B compartment annotation

Insulation scores and annotations for A and B compartments were calculated using cworld (https://github.com/dekkerlab/cworld-dekker). Insulation scores were calculated using contact maps binned at 40 kb resolution, and A and B compartment annotations were calculated using contact maps binned at 200 kb and 1 Mb resolution. Insulation scores were calculated using the script matrix2insulation.pl with the parameters “--ss 80000 --im iqrMean --is 480000 --ids 320000” and compartment annotations were calculated using the script matrix2compartment.pl with default parameters.

#### Detection scores for 3D genome structures

Detection scores were calculated to identify various 3D genome structures in single cells. These structures included chromosome territories, compartments, TADs, centromere interactions, nuclear speckle interactions and nucleolar interactions. Each score reflects how clearly defined a given structure is in a single cell. For example, a clearly defined chromosome territory in a single cell consists of chromosomes interacting more with themselves than with each other (illustration **Figure 2d**). Detection scores were calculated for each structure in each cell as follows:

- Chromosome territories: (observed intra-chromosomal contacts) / (total possible intra-chromosomal contacts) – (observed inter-chromosomal contacts) / (total possible inter-chromosomal contacts)
- Compartments: (observed intra-compartment contacts) / (total possible intra-compartment contacts) – (observed inter-compartment contacts) / (total possible inter-compartment contacts)
- TADs: (observed intra-TAD contacts) / (total possible intra-TAD contacts) – (observed inter-TAD contacts) / (total possible inter-TAD contacts)
- Centromere interactions: (observed centromere-centromere contacts) / (total possible centromere-centromere contacts) – (observed centromere-non-centromere contacts) / (total possible centromere-non-centromere contacts)
- Nuclear speckle interactions: (observed speckle-speckle contacts) / (total possible speckle-speckle contacts) – (observed speckle-non-centromere contacts) / (total possible speckle-non-centromere contacts)
- Nucleolar interactions: (observed nucleolar-nucleolar contacts) / (total possible nucleolar-nucleolar contacts) – (observed nucleolar-non-centromere contacts + observed non-nucleolar-non-nucleolar contacts) / (total possible nucleolar-non-centromere contacts + total possible non-nucleolar-non-nucleolar contacts)

These scores were calculated using a binary contact matrix for each cell, which defined whether or not each pair of genomic bins were in contact in that cell. For TADs, 40-kb bins were used and for all other structures 1-Mb bins were used. Centromere interactions were defined as interactions between positions 3 Mb and 13 Mb of each chromosome. Nuclear speckle interactions and nucleolar interactions were defined as interactions between nuclear speckles regions or nucleolar regions, respectively; the locations of these regions were obtained from Quinodoz et al^7^. To normalize detection scores, an expected detection score was calculated for each 3D genome structure in each cell. The expected detection score was calculated as the mean detection score for 1000 randomized structures, which were generated by randomly shuffling the genomic coordinates of known structures. The normalized detection score for each structure in each cell was calculated as the observed detection score minus the expected detection score.

### Frequencies of higher-order interactions

To determine the percent of cells that contained centromeric or nucleolar interactions, we first calculated an interaction matrix between all 1Mb genomic bins, where the values in the interaction matrix were the percent of cells containing an interaction between each pair of 1Mb bins. We then calculated the mean value for pairs of regions in this interaction matrix representing centromere-proximal or nucleolar regions. For example, to determine the percent of cells containing an interaction between the centromere-proximal regions on chromosome 1 and chromosome 2, we calculated the mean value in this interaction matrix for chromosome 1 positions 3,000,000 to 13,000,000 with chromosome 2 positions 3,000,000 to 13,000,000.

### Higher order structures in scHiC data

Ensemble and single-cell contact maps from scHiC^16^ were plotted to visualize centromere, speckle, and nucleolar interactions. The single-cell barcode from scHiC that was referenced was “hyb_2i-1CDES-1CDES_p10.H9-adj”.

### Identifying heterogeneity in between AB compartments and TADs

Insulation scores, boundary strengths, and compartment scores were generated using *cworld* on the ensemble scSPRITE dataset comprised of the filtered 1000 cells. Insulation scores and boundary strengths were used to identify TADs and compartment scores were used to identify AB compartments in the ensemble scSPRITE dataset. To identify regions of heterogeneity, a genome-wide heatmap was made using the ensemble scSPRITE dataset to look for pseudo-TAD structures in between designated AB compartments and TAD regions. Once a region was identified in a given chromosome, the two 40-kb bins that made up the outermost interaction of the pseudo-TAD structure in the ensemble dataset were used to look for this same interaction in the single-cell dataset (further referred to as “Position A” and “Position B”). For the SE-promoter interaction at the *Nanog* locus, we used the 40-kb bins containing the location of the *Phc1* enhancer and *Nanog* promoter identified previously^31^. In every single cell, we first identified cells containing a contact anywhere along Position A and Position B in that chromosome to ensure that coverage was accounted for. Once the cells with coverage were identified, we identified and grouped cells in this set had contained or lacked the interaction at the intersection of Position A and Position B. **Virtual 4C analysis**

To identify contacts with a specific locus, such as Nanog, we first calculated a contact frequency matrix for all pairs of genomic bins at 40 kb resolution. To convert this contact matrix to a 1-dimensional profile of contacts, we simply used the values in the row of the contact matrix corresponding to the locus of interest.

### Checking for non-cell cycle biases in heterogeneity

We checked the cell cycles on the cells to ensure the differences we were seeing were not the result of biases in cell cycle. We computationally sorted the cells into M, G1, G2, or S phases of cell cycle based on the parameters described previously^16^. After categorizing the cells by phase, we calculated the percentage of cells in each corresponding cell cycle phase in the sets that contained or lacked a particular interaction.

### Human-mouse mixing experiment

To determine the percent of single cells that are mixed together during scSPRITE (from crosslinking until the end of in-nuclei barcoding), we performed a scSPRITE experiment using two different cell types. Mouse embryonic stem cells (bsps) and human cells (HEK293) were mixed together in equal quantities after harvesting both cell lines, and both were crosslinked, digested, dA-tailed, and barcoded in-nuclei together as described previously. Four rounds of in-nuclei barcoding were done (DPM, odd, even, y-even). Nuclei were filtered through a 10-µm filter (PluriStrainer) after in-nuclei barcoding. From the filtered nuclei, 300 nuclei were then reverse-crosslinked and amplified, and libraries were gel-cleaned similarly as described above. Libraries were sequenced using MiSeq, and reads were then aligned to the hg19 and mm9 reference genomes. Reads were sorted into individual cells based on cell-specific barcodes, and we calculated the percentage of reads that aligned to each genome for each identified cell-barcode. We categorized cell-barcodes as mouse- or human-derived when they contained >95% single-species reads, and as mixed when they contained <95% single species reads.

## Detailed Author Contributions

**M.V.A**. Contributed in scSPRITE method development and optimization (Figure 1); Performed scSPRITE experiment to generate data used throughout the paper (Figure 1-5); Performed mouse-human mixing experiment (Figure 1); Generated the ensemble and single cell heatmaps for validation of method + higher order structure (Figure 1, 2, 3, 4, S2, S3, S4); Generated table and plot denoting differences in contacts between scSPRITE and scHiC methods (Figure S2); Identified regions for TAD heterogeneity (Nanog-SE; Tbx3-Lhx5; AB compartment in chr4); generated scripts and plotted heatmaps to demonstrate these differences (Figure 4, 5, S4, S5); Wrote and edited manuscript.

**J.W.J**. Contributed conceptually in scSPRITE method optimization (Figure 1); Designed mouse-human mixing experiment and cultured cells for it (Figure 1); Analysis and plotting: generated cartoon and comparison of contacts and reads between scSPRITE and scHiC (Figure 2); generated histograms & TAD clustering maps using the single cell TAD detection scores (Figure 4, S4); Contributed conceptually to figure design, heatmaps and plots generation (Figure 1-5); Contributed to discussions & biological interpretations (Figure 4, 5, S4, S5); Compiled the figures (Figure 1-5); Wrote and edited manuscript.

**N.O**. Defined and wrote the script for the detection score analysis to quantify single cell genomic structures (Figures 2, 3, 4, S2, S3, S4); Performed the virtual 4C analysis for the plot in (Figure 5;) Performed cworld analysis to generate insulation scores and annotations for A and B compartments; Contributed to writing the manuscript.

**M.S.C**. Contributed in determining optimal crosslinking conditions for scSPRITE (inferred in Fig 1); Contributed to optimizing in-nuclei biochemical processes(nuclei isolation, in-nuclei digestion & dA-tailing) (Figure 1); Major contributor in developing the scSPRITE in-nuclei barcoding workflow and nuclei sonication conditions (Figure 1).

**C.A.L**. Wrote the pipeline to identify cell-specific barcodes and to group complexes containing the same cell-specific barcode from sequencing data.

**S.A.Q**. Contributed to experiments for scSPRITE method development and validation (Figure 1).

**D.A.S**. Conceptualized the idea for scSPRITE; Contributed to scSPRITE method development (nuclei isolation, in-nuclei digestion & dA-tailing, in-nuclei barcoding workflow) (Figure 1).

## ACKNOWLEDGEMENTS

We would like to thank the assistance from Fan Gao from Caltech’s Bioinformatics Resource Center and Igor Antoshechkin from Caltech’s Millard and Muriel Jacobs Genetics and Genomics Laboratory. We would like to thank Chris Chen, Vicky Trinh, Elizabeth Detmar, Elizabeth Soehalim, and Aditi Narayanan for their contributions in helping develop scSPRITE. We would like to thank Matt Thompson’s lab for allowing us to use their MiSeq instrument. We also thank Natasha Shelby and Shawna Hiley for contributions to writing and editing this manuscript and Inna-Marie Strazhnik for helping with illustrations.

## AUTHOR CONTRIBUTIONS

M.V.A. conducted the experiments to develop and validate method, conceptualized and performed the analyses, wrote the manuscript; J.W.J contributed the experiments to develop and validate the method, conceptualized and performed analyses, wrote the manuscript; N.O conceptualized and performed analysis to validate the method, developed the pipeline for the workup of scSPRITE sequencing data, contributed to writing the manuscript; M.S.C contributed to the experiments to develop the method; C.A.L developed a pipeline to sort cells by cell-specific barcodes; S.A.Q contributed to the experiments to develop and validate the method; D.A.S contributed to conceptualize scSPRITE and to the experiments to develop the method; M.G. conceptualized scSPRITE, supervised the experiments and the analysis to validate the method, wrote the manuscript; R.F.I conceptualized scSPRITE, supervised the experiments and analysis to develop the method, wrote the manuscript.

## DATA AVAILABILITY STATEMENT

The datasets (Fig. 1-5; supplemental figures S1-S5) generated during and analyzed during the current study are available in the GEO repository under accession number GSE154353 (https://www.ncbi.nlm.nih.gov/geo/query/acc.cgi?acc=GSE154353). scSPRITE software is available at https://github.com/caltech-bioinformatics-resource-center/Guttman_Ismagilov_Labs.

---

**Figure S1:**
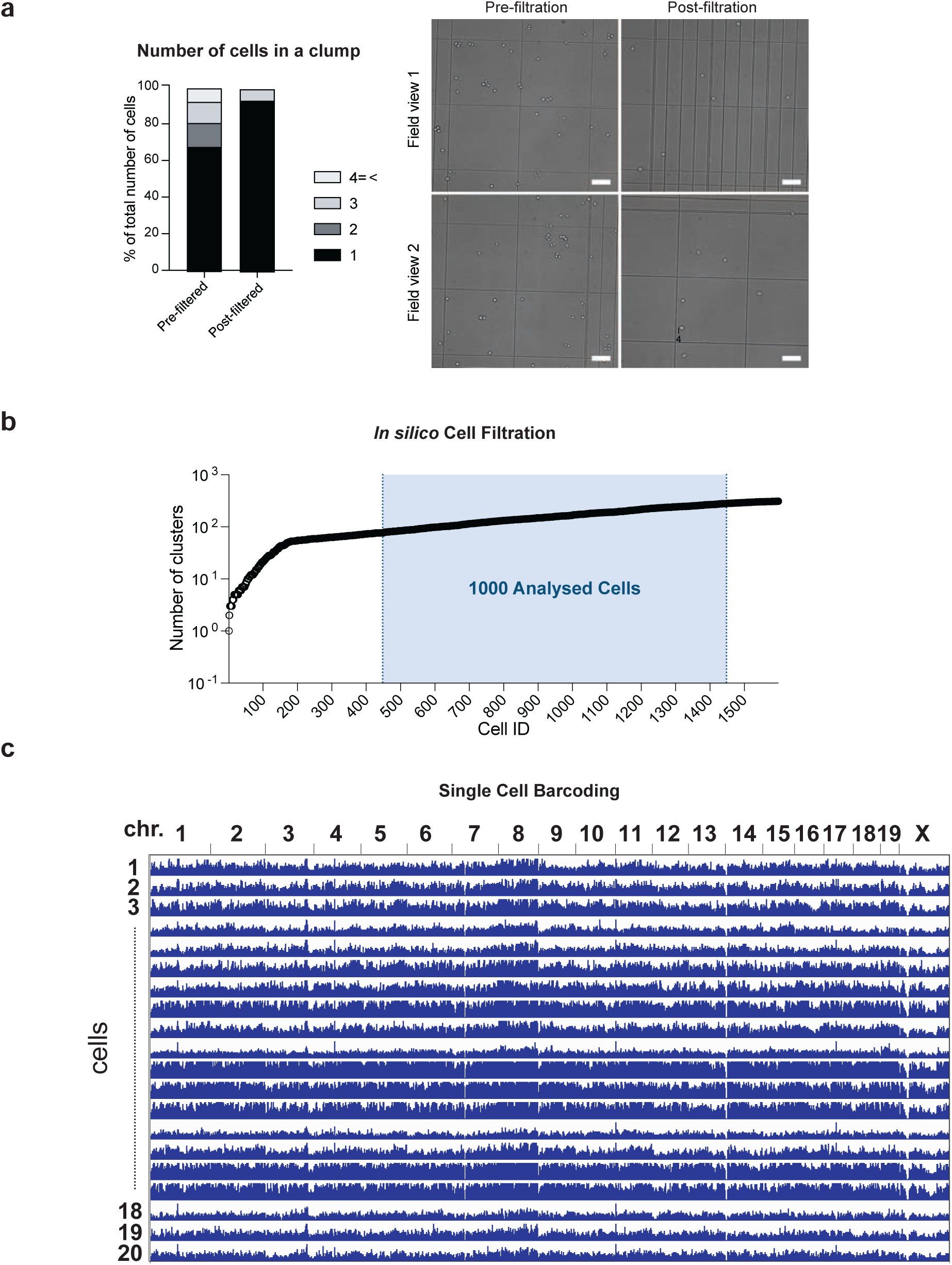
Single cell SPRITE protocol provides single cells with high genomic coverage. **a**, Quantification of cell clumping: on the right microscope images (10x) of cells pre- and post-filtration step, on the left plotted number of cells in clumps pre- and post-filtration (singlets, doublets, triplets, etc); scale bar 100um. **b**, In silico filtration steps: first the top 3.4% of cell barcode IDs containing the largest number of DNA clusters (DNA fragments with same DNA barcode and in-nuclei barcode) are removed (based on the estimation from Figure 1b), then cell barcode IDs with low number of DNA clusters are removed; only 1000 cells (indicated in green) were further analyzed. **c**, Genomic coverage of 20 representative cells barcoded and analyzed with scSPRITE workflow; 1Mb bin per chromosome.

**Figure S2:**
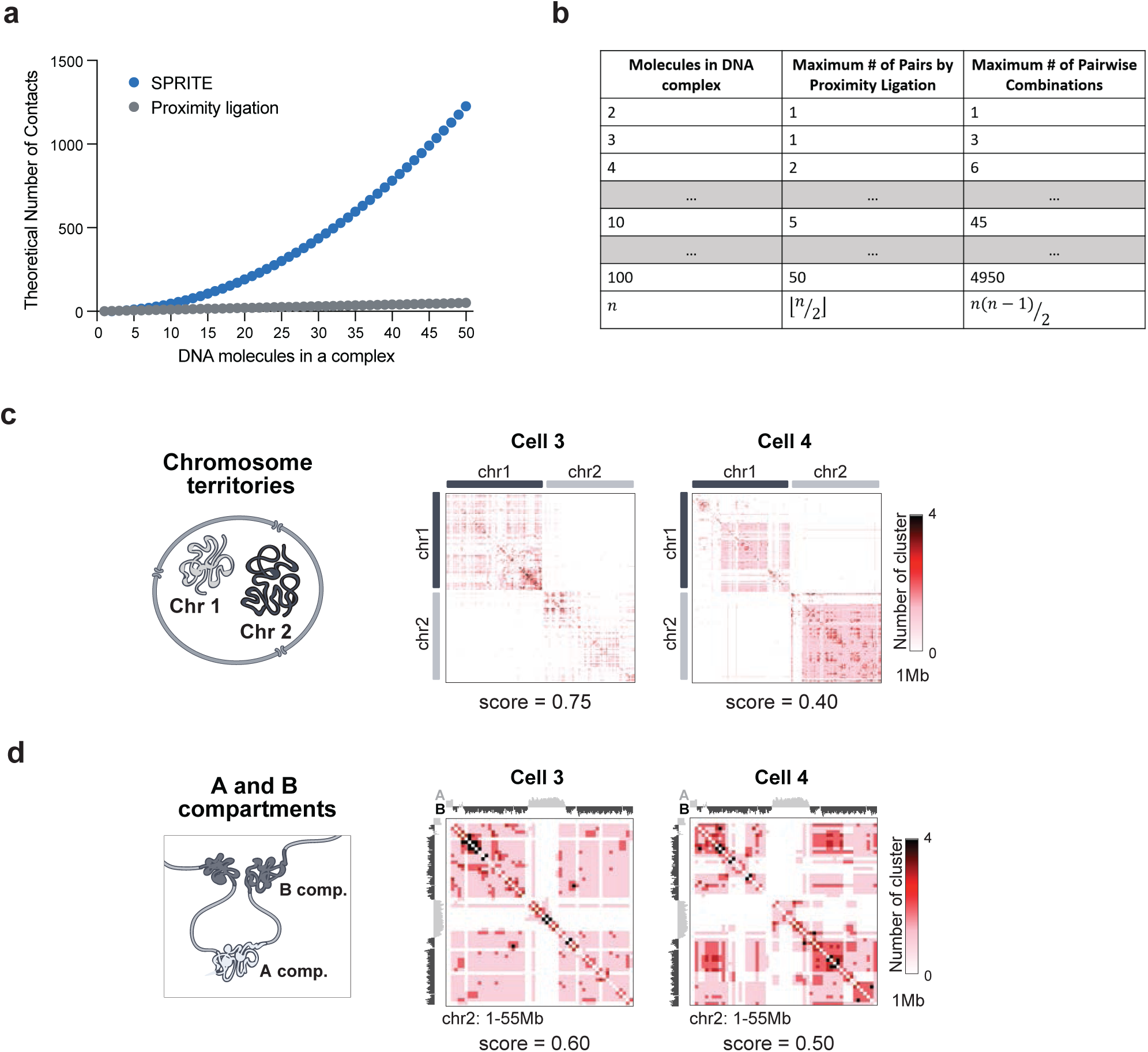
Differences in measurement of multiway contacts versus pairwise contacts. **a**, Plot depicting the theoretical number of contacts measured by SPRITE-derived methods and HiC-derived methods over increasing numbers of DNA molecules per complex. **b**, Comparison of the maximum number of pairwise interactions that can be obtained from proximity ligation (HiC-derived methods) vs the maximum number of pairwise interactions possible over increasing numbers of DNA molecules per complex (SPRITE-derived methods). **c**, Single cell examples of chromosome territory structure between chr1 and chr2; contact map represents number of DNA clusters at 1Mb resolution; detection scores below contact map. **d**, Single cell examples of A/B compartments detected within 0-55Mb in chr2; contact map represents number of DNA clusters at 1Mb resolution; detection scores below contact map

**Figure S3:**
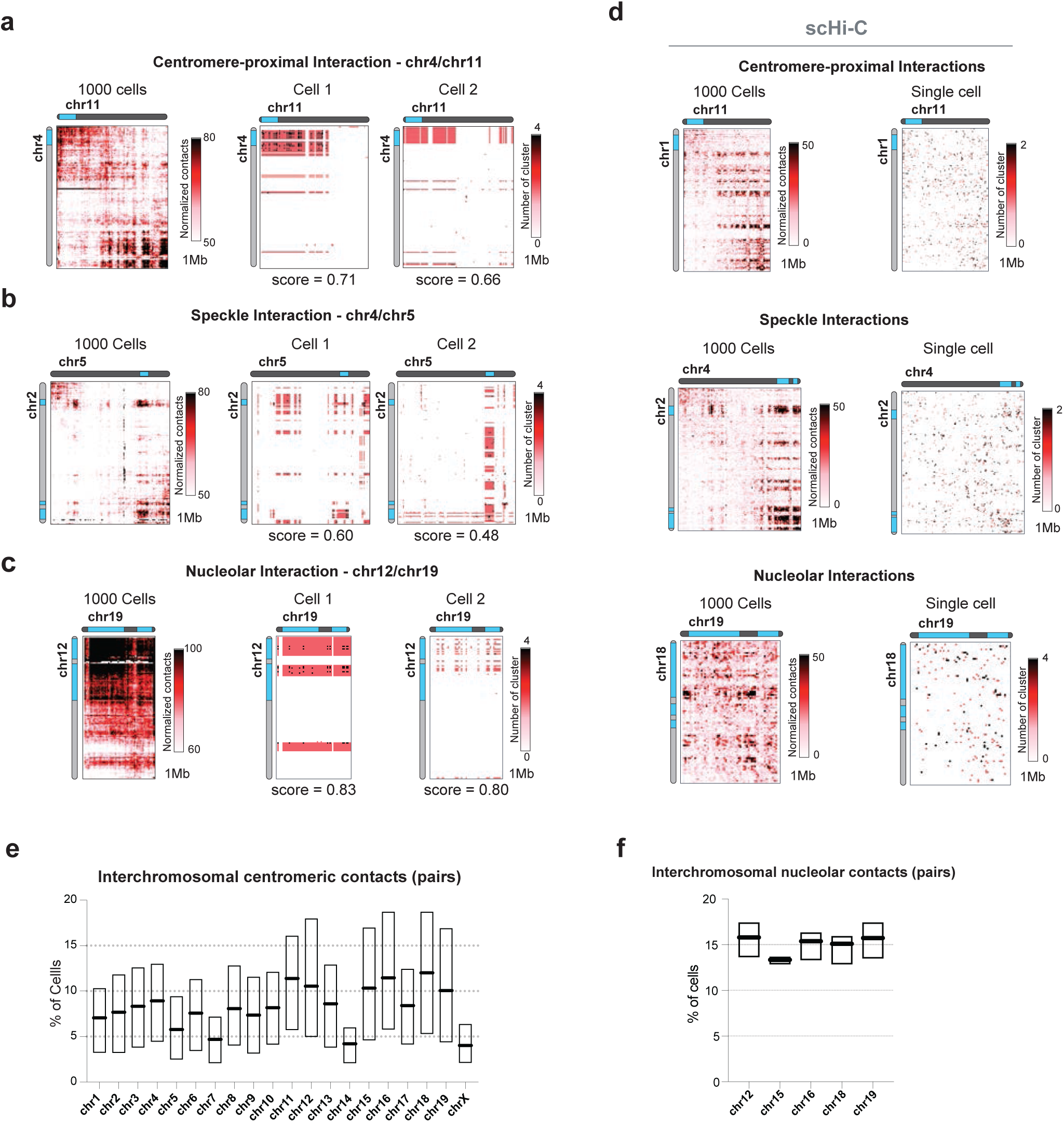
Higher-order structures are identified genome-wide in hundreds of single mESC by scSPRITE method. **a**, Single cell examples of chr4 and chr11 centromere-proximal regions interacting together; contact map represents number of DNA clusters at 1Mb resolution. **b**, Single cell examples of speckle interaction detected between chr2and chr5; contact map represents number of DNA clusters at 1Mb resolution. **c**, Single cell examples of nucleolar interactions detected between chr12 and chr19; contact map represents number of DNA clusters at 1Mb resolution. **d**, Higher-order structures representation from scHiC data16 – centromere-proximal interactions, speckle interactions, and nucleolar interactions; Pairwise contact map from ensemble 1000 cells (left), pairwise contact map from their best single cell (right). **e**, Barplot representing the percent of cells containing interchromosomal centromere contacts with another chromosome for each chromosome in the genome. Bolded black bar represents the mean. **f**, Barplot representing the percent of cells containing interchromosomal nucleolar contacts with another chromosome for each nucleolar associating chromosome. Bolded black bar represents the mean.

**Figure S4:**
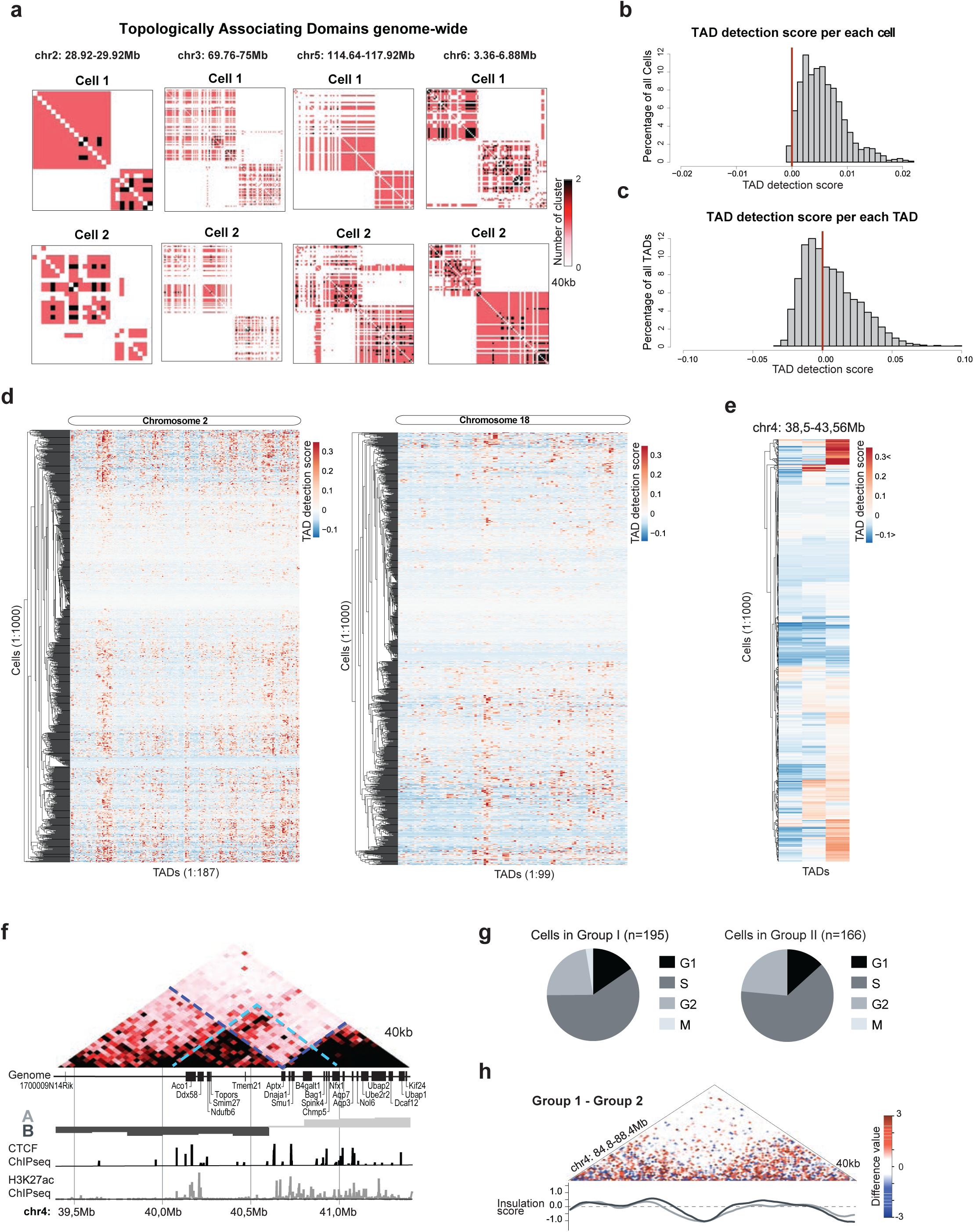
TADs are heterogeneous units present in the genomes of individual mESCs. **a**, Examples of contact map demonstrating TAD-like structures in single cells across different chromosomes; 40kb resolution. **b**, Histogram representing normalized detection scores across all 1000 cells per each TAD detected in ensemble scSPRITE data; red line marks score=0 above which detected structure is similar to the contact map where all potential contacts within a TAD structure are made. **c**, Histogram representing TAD detection scores across all TADs detected in ensemble scSPRITE data per single cell; red line marks score=0 above which detected structure is similar to the contact map where all potential contacts within a TAD structure are made. **d**, Representation of TAD detection scores across 1000 cells in chr2 (left) and chr18 (right). Columns represent the strength of TAD detection scores for all TADs detected across ch2 or chr18 respectively in ensemble scSPRITE; rows represent each single cell from 1000 analyzed cells clustered based on score similarity pattern. **e**, Representation of TAD detection scores across 1000 cells between 38.5-48.56Mb of chr4. Each line represents strength of TAD detection scores in this given region from a single cell. **f**, Ensemble heatmap from all 1000 cells between 39.4-41.4Mb of chr4 representing strong TADs detected in bulk (blue lines), and weak emerging TAD (green line) over the A/B boundary. Corresponding genome tracts and ChIPseq tracks for CTCF and H3K27ac are presented below. **g**,Pie chart representing the fraction of cells in each cell cycle phase from the set of single cells containing (left) or lacking (right) the contact between the boundary region (Figure 4e). **h**, Difference contact map across a control region 84.8-88.4 Mb of chr4 made by subtracting the normalized contacts from cells in Group II from Group I (Figure 4d). Insulation scores for cells in Group I (dark grey) and Group II (light grey) are plotted.

**Figure S5:**
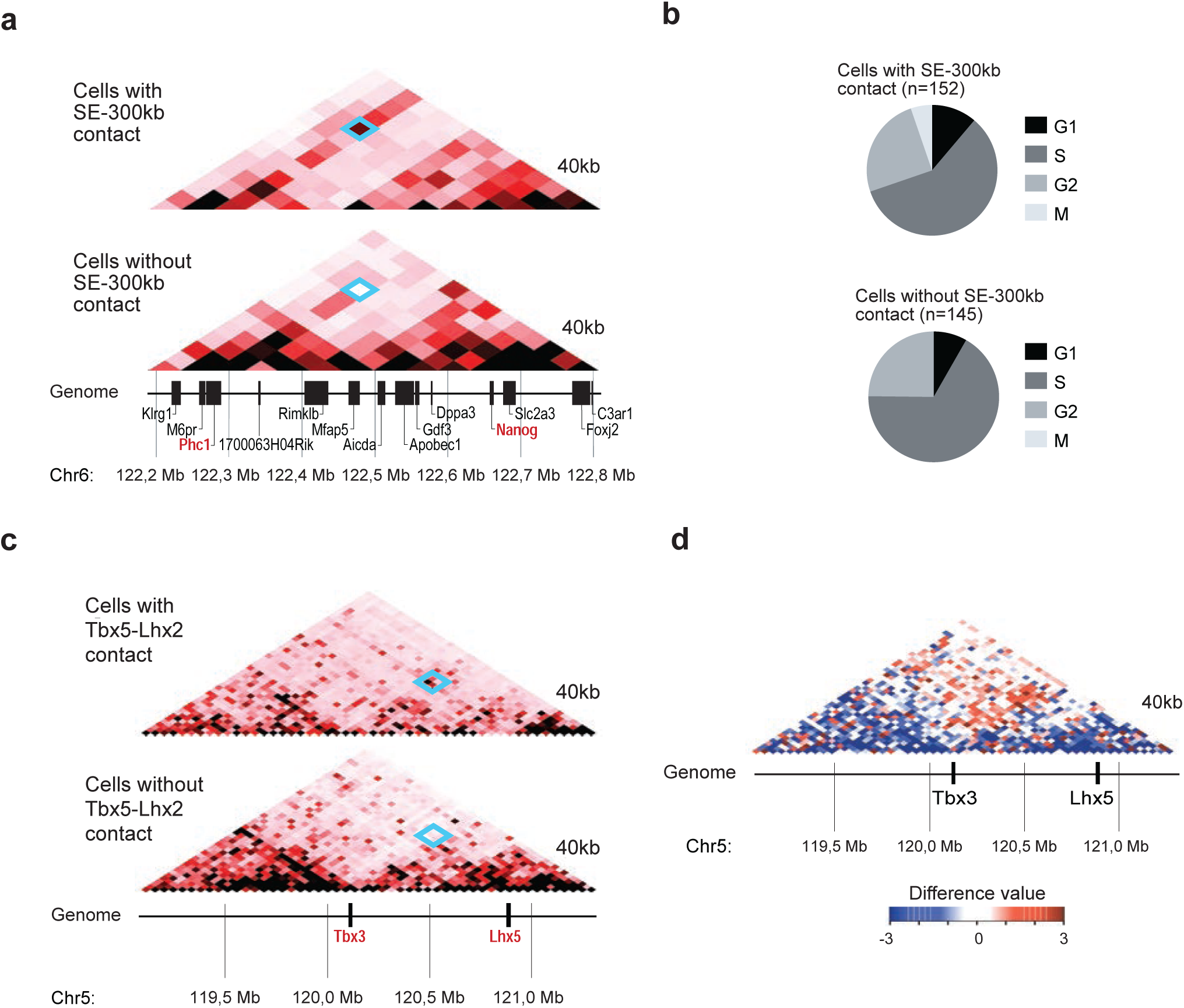
Structural heterogeneity in promoter-enhancer interactions is revealed by scSPRITE. **a**, Ensemble heatmaps across 122.2-122.8Mb region in chr6 representing cells containing (top) or lacking (bottom) the contact between the Nanog locus and the -300 Kb SE. Blue square demarcates the contact. **b**, Pie chart representing the fraction of cells in each cell cycle phase from the set of single cells containing (left) or lacking (right) the contact between the Nanog locus the SE 300kb upstream of Nanog. **c**, Heatmaps between 119.24-121.28Mb in chr5 of pooled cells either containing (top) or lacking (bottom) the contact between the Lhx5 locus and the Tbx3 enhancer. Blue square demarcates the contact. **d**, Contact map across 119.24-121.28 Mb of chr5 highlighting regions that are enriched or depleted of the contact from Figure S5c.

